# Measures of reproducibility in sampling and laboratory processing methods in high-throughput microbiome analysis

**DOI:** 10.1101/322677

**Authors:** Patricia Vera-Wolf, Juan P. Cárdenas, Amanda M. Morton, Tomás Norambuena, Rafael Torres, Luis E. Leon, Elisabeth M. Bik, Juan A. Ugalde, Daniel E. Almonacid, Jessica Richman, Zachary S Apte

## Abstract

Microbial community analysis can be biased by multiple technical factors, such as storage conditions, DNA extraction, or amplification conditions. In a high-throughput laboratory that relies on samples obtained from thousands of different subjects, knowledge of the extent of subject-introduced sampling and storage variation on the outcome of the inferred microbiome, as well as the effect of laboratory-introduced variation caused by reagent batches, equipment, or operator on the consistency of these processes within the laboratory is paramount. Here, we analyzed the effect of sampling from different parts of the same stool specimen or on different consecutive days, as well as short-term storage of samples at different temperatures on microbiome profiles obtained by 16S rRNA gene amplification. Each of these factors had relatively little effect on the microbial composition. In addition, replicate amplification of 44 stool samples showed reproducible results. Finally, 363 independent replicate extractions and amplifications of a single human homogenized stool (HS) specimen showed reproducible results (average Lin’s correlation = 0.95), with little variation introduced by HS batch, operator, extraction equipment, or DNA sequencer. In all cases, variations between replicates were significantly smaller than those between individual samples; subject identity always was the largest determinant. We propose that homogenized stool specimens could be used as quality control to routinely monitor the laboratory process and to validate new methods.

## Introduction

The necessity of standardization in experimental microbiome analysis is a relevant aspect for its implementation in human research, as well as for clinical approaches. Among the components required for standardization, reproducibility is one of the most important aspects (Nature Microbiology Editorial 2016). Microbiome analysis involves the analysis of the composition of the microbial communities found within a biological sample, such as feces or soil. Since microbiome studies generally aim to analyze the composition of a complex microbial community, the goal is to recover the presence of each microbial species within a mixture with equal chance. Ideally, the microbiome composition as found after processing in the lab should match that of the original, unprocessed sample. However, individual microbial species vary from each other in their abilities to grow under suboptimal storage conditions, to be lysed during DNA extraction, or to be amplified with marker gene PCR methods (Bag *et al.* 2016). Therefore, each choice of method during storage, DNA extraction and amplification steps will be more efficient for certain microbial taxa than for other groups, and thus affect the outcome - almost no microbiome analysis can exactly reproduce the composition of the original biological sample (Brooks *et al.* 2015; de la Cuesta-Zuluaga and Escobar 2016; Costea *et al.* 2017). Because each alternative processing method can eventually lead to different microbiome profiles, it is challenging to compare microbiome studies across studies and research groups (Tyler *et al.* 2014; Bik 2016; Costea *et al.* 2017 Kim *et al.* 2017; Pollock *et al.* 2018).

Storage conditions are a first set of variables that have been shown to affect microbiome analysis outcomes (Röder *et al.* 2010; Choo *et al.* 2015; Romanazzi *et al.* 2015). Ideally, a biological sample taken for microbiome analysis should be freshly analyzed (Pollock, *et al.* 2018) or stored at very cold temperatures, such as −80°C, immediately after sampling to preserve the microbiological composition by preventing selective growth of microorganisms (Vandeputte *et al.* 2017). However, immediate cryostorage is often not feasible, for example in field conditions without reliable access to electricity (Schnorr *et al.* 2014; Hale *et al.* 2015; Vogtmann, Chen, Kibriya, *et al.* 2017) or for subjects who obtain specimens in their own homes and send them to a laboratory per regular mail (Schultze *et al.* 2014; Fu *et al.* 2016; Al *et al.* 2018). Fecal samples have been shown to be relatively stable at room temperature for short periods of time (Carroll *et al.* 2012; Tedjo *et al.* 2015; Guo *et al.* 2016; Bassis *et al.* 2017; Wong *et al.* 2017), but storage and shipping of samples without preservatives at room temperature for periods longer than 24 h has been shown to affect the microbial composition (Sinha *et al.* 2016; Song *et al.* 2016; Amir *et al.* 2017). Several methods have been developed to preserve microbial composition in cases that immediate storage at below-freezing temperatures is not feasible. These include storage on fecal cards, storage in ethanol, or stabilization buffers (Anderson *et al.* 2016; Hill *et al.* 2016; Song *et al.* 2016; Vogtman, Chen, Kibriya *et al.* 2017; Vogtmann, Chen, Amir *et al.* 2017; Vandeputte 2017).

The choice of DNA extraction technique is another factor that can greatly influence the outcome of microbial analysis. The most popular commercial DNA extraction methods are all based on guanidine-containing buffers and silica gel spin columns (Thatcher 2015; Thomas *et al.* 2015). However, variations between these kits, and the additional choices of inclusion of bead beating, lysozyme, Proteinase K, and other additives vary widely between suppliers and laboratories and have all been shown to have an influence on the microbiome analysis (Kennedy *et al.* 2014; Brooks *et al.* 2015; de la Cuesta-Zuluaga and Escobar 2016; Gerasimidis *et al.* 2016; Hiergeist *et al.* 2016; Knudsen *et al.* 2016). For example, bead beating has been shown to affect the inferred microbial composition of biological replicates, because it increases the lysis of gram-positive bacteria (Yuan *et al.* 2012; Knudsen *et al.* 2016; Pollock *et al.* 2018).

After DNA extraction, there are several other technical variables that could lead to biases in taxonomic profiles. Many microbiome studies rely on the amplification and sequencing of marker genes, most commonly the 16S rRNA gene (Knight *et al.* 2017; Young 2017). The choice of PCR primers and amplification conditions has been shown to be an important source of bias and variation in microbiome studies (Klindworth *et al.* 2013; Walker *et al.* 2015; de la Cuesta-Zuluaga and Escobar 2016; Fischer *et al.* 2016; Gohl *et al.* 2016; Thijs *et al.* 2017). For both amplicon based as well as shotgun based microbiome profiling, the choice of sequencing library kit (Jones *et al.* 2015) or bioinformatics pipeline (Allali *et al.* 2017) can have an influence on the inferred taxonomy. In addition, taxonomic profiles can also vary between 16S rRNA gene amplification and shotgun sequencing (Jovel *et al.* 2016; Ranjan *et al.* 2016; Tessler *et al.* 2017).

For a high-throughput laboratory that follows the same extraction and amplification standard operating procedures for thousands of samples, the effect of other technical and human variables on the observed microbiome profiles are of interest as well. Some of these variations are within the control of the laboratory and can also be tracked, such as variations between reagent batches, equipment, or operators. Another set of technical variations could be introduced by the subject providing the sample, such as sampling different parts of a fecal specimen, sampling on consecutive days, or storing a sample at a different temperatures. Because these variables are introduced before a sample reaches the laboratory, they fall outside the quality control measures of a laboratory, and their effects on the outcome of microbial analysis are less well understood. However, in the setting of a laboratory obtaining samples from thousands of different individuals, each of whom might have a different sampling or storage approach, it is important to study their contribution to variations in microbiome profiles.

Here, we address the relative influence of several of user and laboratory-based variations. We first compared the effect of sampling different parts of the same stool specimen as well as the effect of sampling on nearly adjacent days. Secondly, the effect of storage at several temperatures of samples in a lysis and stabilization buffer was investigated. Thirdly, we determined how 3 independent rounds of amplification and sequencing performed on the same DNA affected the composition of the microbiome. Finally, we compared the microbiome profiles obtained from 363 replicate aliquots of the same homogenized stool sample, extracted, amplified, and sequenced with an identical protocol.

## Methods

### Informed consent and IRB approval

This study was approved under a Human Subjects Protocol provided by an independent IRB (E&I Review Services, IRB Study #13044, 05/10/2013). E&I is fully accredited by the Association for the Accreditation of Human Research Protection Programs, Inc. (AAHRPP), with Registration # IRB 00007807. uBiome undergoes a yearly voluntary continuing review by E&I Review Services to determine that the project meets the same scrutiny of human subjects protections as projects conducted at research institutions. The IRB board membership of E&I Review Services is consistent with the Code of Federal Regulations (CFR) requirements of 21 CFR 56.107 and 45 CFR 46.107. Informed consent for participation in this study was obtained from all subjects. All participants were 18 years or older.

### Overview of self-sampling, shipping, and microbiome analysis

uBiome has developed and protected a method for the analysis of the microbiome obtained by an individual (Apte and Richman 2015). This patent covers the sampling kit contents, the methods to obtain biological samples by self-sampling (e.g. stool samples can be obtained from toilet paper using a sterile swab), transport and storage of specimens at room temperature in a lysis and stabilization (LS) buffer, the generation of a microbiome sequence dataset, and the transmission of analyzed data to the individual (Apte and Richman 2015). In addition, uBiome has protected technology (U.S. granted patent) to analyze the microbiome composition for a population of subjects and to use this to generate a functional diversity dataset (Apte *et al.* 2017); based on this patent, we identified a set of taxa associated with gastrointestinal conditions, which was described previously (Almonacid *et al.* 2017).

### Duplicate sampling from stool in a 20-day period

A healthy male subject, subject A, in his thirties, collected duplicate stool samples over a period of 20 days, during which he maintained his regular, omnivorous diet. During each of 11 stool collection events, two distinct pieces of fecal material spaced 1 to 4 cm apart were collected from the same piece of toilet paper after defecation, each using a separate sterile swab. Swabs were swirled for one minute, each in a separate uBiome vial containing LS buffer and zirconia beads. Samples were stored at room temperature for a maximum of 3 weeks and extracted as described below.

### Stability at different temperatures

Eight different donors (subjects 1 through 8) collected stool specimens on the same day. Each subject sampled 10 pieces of stool from the same piece of toilet paper into 10 different vials with 750 ul of LS buffer. The 10 specimens per person then were pooled and homogenized by vortexing, to obtain 8 different pools of 7.5 ml each. Each homogenized stool sample then was used to make 10 aliquots of 500 ul each. These 10 aliquots (labeled A through J) were used to test 5 different storage conditions, with two replicates per storage condition. Replicates A and B were immediately used for DNA extraction and the DNA was stored at 4°C for a week. Replicates C through J were stored for a week at different temperatures as follows: at −20°C (C and D), at 20°C (E and F), at 30°C (G and H), or at 40°C (I and J), after which samples were processed by DNA extraction. The extracted DNA from all 80 replicates then were used for 16S rRNA gene PCR amplification and sequencing as described below.

### Replicate amplification of 44 stool samples

A set of 44 stool samples, each from a different subject, were used to check the effect of replicate 16S rRNA gene amplification and sequencing runs performed on the same extracted DNA. After DNA extraction, the 44 samples were amplified in triplicate and sequenced in three independent runs to yield a total of 132 microbiome profiles (see below).

### Replicate extractions and amplifications of a single homogenized stool

A single complete stool specimen was self-collected by a healthy male donor, donor B (a different donor than person A, 30 years old). Immediately after the defecation event, the complete bowel movement (85.3 grams) was mixed with 1000 ml LS buffer and homogenized in a glass kitchen blender (Oster) for 10 minutes at low speed, followed by 10 min at high speed. The sample was then aliquoted in 50 ml portions in sterile falcon tubes, and stored frozen at −20°C. To prepare a new batch for use, a single 50 ml aliquot was taken from the freezer and thawed overnight in a fridge and mixed subsequently by vigorously shaking by hand. Then, 10 ml of this stock was mixed with 30 ml of LS buffer and mixed on a vortexer for 6 minutes. The mixture was then left at room temperature for 30 to 60 min to allow for settlement of larger particles, and the supernatant was pipetted into a new 50 ml falcon tube and 50 ul portions was added into uBiome tubes with LS buffer and zirconia beads. These homogenized stool (HS) aliquots were kept at 4°C and used within 3 months. For this study, 363 HS samples prepared in 4 different batches, but all derived from the same original homogenized stool specimen, were analyzed. Each of these replicates was processed in a different, independent DNA extraction and amplification run (one HS replicate per set of 96 samples).

### Other human stool samples

From a set of stool samples from 897 healthy subjects described before (Almonacid *et al.* 2017), 400 samples were randomly selected using the R package “sample”, and their datasets were used in this study.

### DNA extraction and amplification

All stool samples were processed using the same DNA extraction, amplification, and sequencing protocols as described before (Almonacid *et al.* 2017). Briefly, stool samples were subjected to bead-beating and DNA extraction, and the V4 variable region of the 16S rRNA gene was amplified with broad range primers, and sequenced. Sequencing was performed in a paired-end modality on the Illumina NextSeq 500 platform rendering 2 × 150 bp paired-end sequences. Sample handling, DNA extraction, amplification, and sequencing was performed using Standard Operating Procedures in a Clinical Laboratory Improvement Amendments (CLIA) licensed and College of American Pathologists (CAP) accredited laboratory. DNA extraction was performed by one individual from a group of 15 rotating operators, on one of 9 DNA extraction robots, and analyzed on one of 3 sequencing machines.

### 16S rRNA sequencing analysis

The bioinformatics analysis was performed as described previously (Almonacid *et al.* 2017). Briefly, forward and reversed 16S rRNA gene V4 sequence reads were demultiplexed, quality-filtered, and merged after primer removal. Only samples with at least 10,000 high-quality reads were used in the analysis. Chimeras were removed using VSEARCH (Rognes *et al.* 2016), and then clustered using Swarm (Mahé *et al.* 2014) . The resulting clusters were then compared to a curated version of the SILVA 123 database (Quast *et al.* 2013), using 100% identity over 100% of the length. The relative abundance of a set of 28 species and genus-level microbial taxa (named “clinical taxa” in this study) known to be associated with health conditions and that could be identified with high precision and sensitivity (Almonacid *et al.* 2017) was determined by dividing the count linked to those taxa by the total number of filtered reads.

Beta-diversity was investigated using Bray-Curtis dissimilarity and non-metric multidimensional scaling (NMDS) ordination as implemented in the R package phyloseq (McMurdie and Holmes 2013). To compare similarity of microbiome profiles to each other, Lin’s concordance correlation coefficient (CCC) of the genus-level clinical taxa was calculated for each community pair (Lin 1989). Statistical significance was tested using the non-parametric Kolmogorov-Smirnov (KS) test (Massey 1951). Permanova analysis was performed on the distance matrix used for beta-diversity analysis using the R Vegan package. (Oksanen *et al.* 2016). Permanova was performed using the function *adonis*, with 999 permutations. Heterogeneous dispersion of the data was evaluated using the function *betadisper*.

To compare the microbiome profiles of the 44 triplicate amplifications, cluster analysis on Bray-Curtis dissimilarity matrices was performed using the Ward’s method (Ward 1963). Tree topology was visualized using the Interactive tree of life (iTOL) software (Letunic and Bork 2016).

## Results

### Duplicate stool sampling on consecutive days

To determine the variation of stool microbial composition associated with taking multiple samples from the same piece of toilet paper, and on (nearly) consecutive days, a single person, subject A, collected 11 paired stool sample over a period of 20 days. The paired stool samples, taken 1 to 4 cm apart from the same piece of toilet paper after elimination, showed very similar microbiome profiles, which was confirmed by an average within-pair Lin’s correlation factor of 0.95 (+/− 0.04) (Figure 1A). The average Lin’s correlation between stool samples collected on different days was 0.68 (+/− 0.17), which was significantly lower than that of the within-pairs (KS test p<0.0001), with no apparent drift over time (Figure 1A). Beta diversity analysis showed that all 22 samples from subject A looked very similar to each other in a comparison with replicate stool samples from 8 different subjects, with a clear clustering by individual (Figure 1B). Subject A’s samples showed clustering of paired samples, again, with no apparent drift over time (Figure 1C).

**Figure 1.**
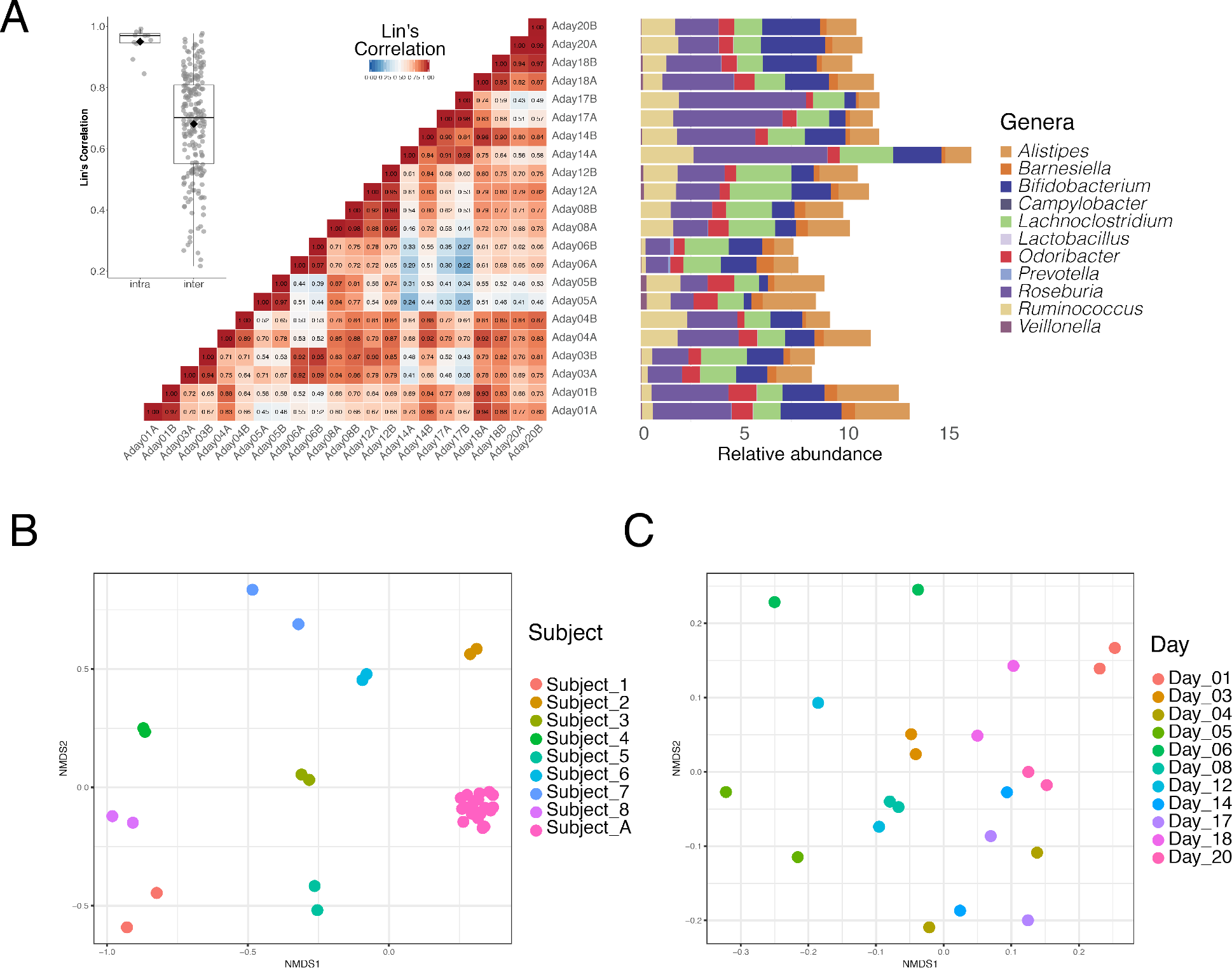
Reproducibility of duplicate sampling from stool and stability of gut microbiome over a 20-day period. Subject A took 11 paired stool samples within a 20-day period. On each sampling day, subject A collected 2 distinct pieces of stool from the same piece of toilet paper. **A.** Heatmap of the Lin’s correlation between samples obtained from the same piece of toilet paper (within-pair) or on different days (between-pairs). The relative abundance of clinical microbial taxa found within the stool samples is plotted as a horizontal bar graph on the right. Samples are labeled per day (e.g., there was no sampling event on day 2). The inset on the left shows a boxplot of Lin’s correlation of within pair and between pair comparisons. **B.** NMDS ordination of Bray-Curtis dissimilarity values of all 22 stool samples from subject A, shown in pink, and replicate stool samples from 8 other subjects, shown in other colors. **C.** NMDS ordination of Bray-Curtis dissimilarity values of the 11 paired stool samples from subject A only.

### Stability of stool samples in a stabilization buffer at different temperatures

To study the effect of short-term storage of stool samples in a lysis and stabilization (LS) buffer at different temperatures, homogenized fecal specimens from 8 different subjects in LS buffer were used for DNA extraction, either immediately after homogenization, or after storage for a week at four different temperatures. Each extraction was performed in duplicate. Samples were compared to each other by plotting the relative abundances of the clinical taxa after immediate extraction against those obtained after storage at different temperatures (Figure 2A and B). The average within-individual Lin’s correlation between the 10 replicate samples from the same subject was 0.92 (SD=0.105). In contrast, the between-person Lin’s correlation between replicate samples from different subjects was 0.28 (SD=0.24); significantly lower than the within-individual correlation (KS test, p<0.0001). In addition, beta-diversity analysis showed no apparent effect on sample composition. Samples clustered much more per subject than per storage condition (Figure 2B and C). Permanova (Adonis) showed no significant differences using temperature as the clustering group (p=0.935) while clustering using subject as the determinant was significant (p = 0.001). In addition, these results are not an artifact of heterogeneous dispersion of the data, based on the betadisper results (p = 0.44).

**Figure 2.**
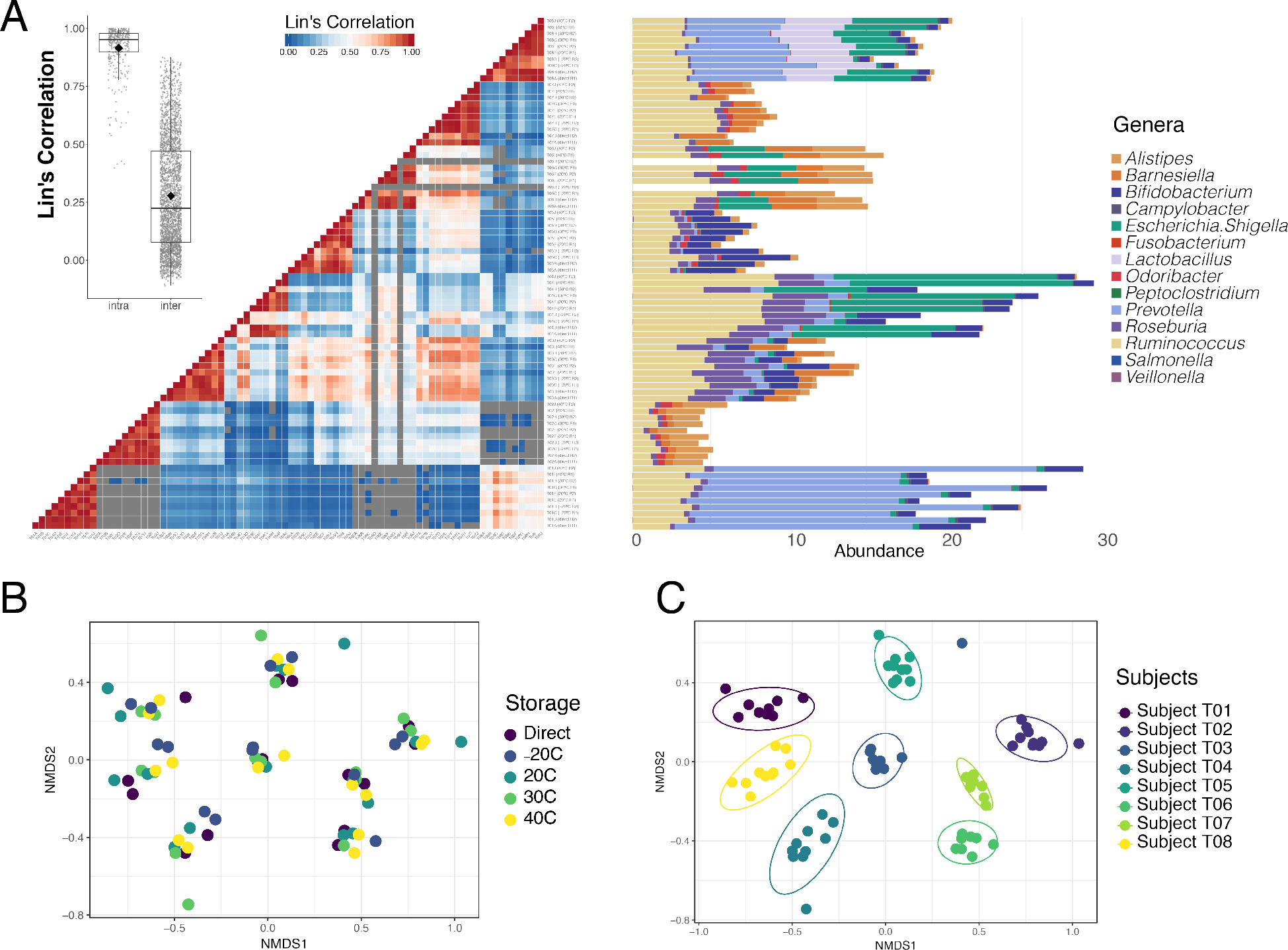
Effect of storage at different temperatures on stool samples from 8 different subjects. Each stool sample was homogenized in LS buffer, then split into 10 different aliquots from which DNA was either extracted directly (and then stored for a week at 4°C), or after a week of storage at four different temperatures (−20°C to +40°C). Each of the 5 storage conditions was tested in duplicate. **A.** Heatmap of the Lin’s correlation between the 10 replicates. Replicates are sorted first per subject (n=8), then per storage condition (n=5), then per replicate (n=2). Negative or missing correlation values are shown in grey. The inset on the left shows a boxplot of Lin’s correlation of within pair and between pair comparisons. The relative abundance of clinical microbial taxa found within the samples is plotted as a horizontal bar graph on the right. **B.** NMDS ordination of Bray Curtis dissimilarity matrix of microbial communities from these samples, showing data-points colored by storage temperature. **C.** Same data points as in D, but colored by original stool sample. Circles indicate clustering with 95% confidence interval.

### Inter-run amplification and sequencing variation in 44 stools

To assess the degree of variation introduced by amplification and sequencing samples, the 16S rRNA gene of extracted DNA from a set of 44 stool samples was amplified and sequenced in triplicate in three independent experiments. The average intra-individual Lin’s correlation between the 3 replicate pairs was 0.94 (SD: 0.12), while the average inter-individual correlation was 0.28 (SD: 0.31) (Figure 3A). This difference was statistically significant (KS test; p-value <0.0001). Beta-diversity analysis showed clustering of the three triplicates of most samples together (Figure 3B); this clustering was confirmed in a radial tree topology (Figure 3C).

**Figure 3.**
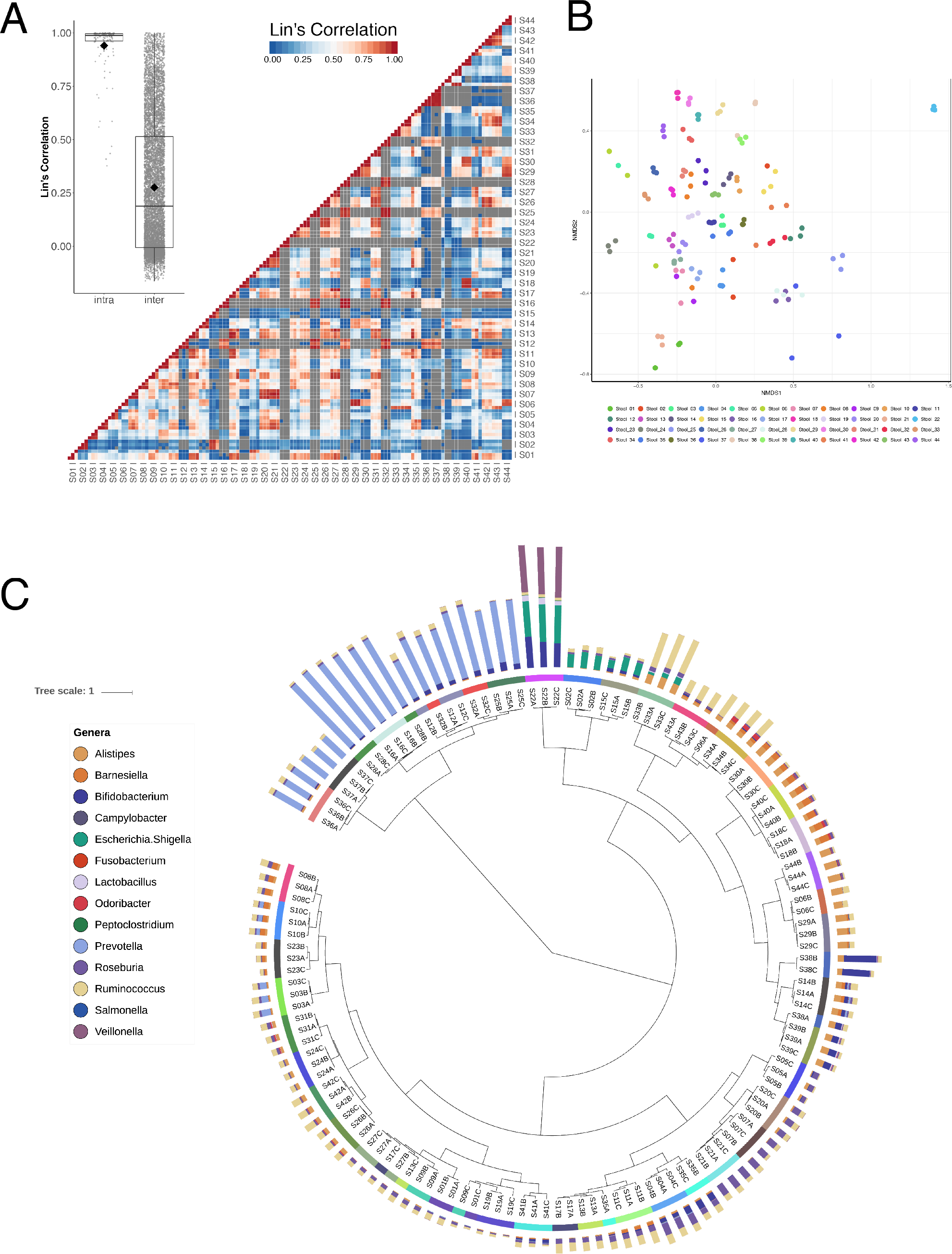
Inter-run amplification and sequence variability of 44 stool samples. Extracted DNA of 44 stool samples was amplified and sequenced in triplicate, in three independent runs. **A.** Heatmap of the Lin’s correlation between all 132 replicates. Replicates are sorted first per subject (n=44), then per replicate (n=3). The inset on the left shows a boxplot of Lin’s correlation of within pair and between pair comparisons. **B.** Beta diversity analysis using NMDS ordination of Bray-Curtis dissimilarity of the replicate amplifications. Stool samples are colored per subject. **C.** Radial dendrogram of the 44 stool samples amplified and sequenced in triplicate. The middle circle shows the identity of the 44 stool samples; each stool sample has a different color. Genus level bars are shown on the outer circle; colors are explained on the key on the left. The tree topology was created using cluster analysis on Bray Curtis dissimilarity matrices using the Ward’s method, and visualized using iTOL (Letunic and Bork 2016).

### Reproducibility of microbiome profiles of 363 replicate extractions

To test the reproducibility of DNA extraction, and amplification methods in a high throughput laboratory setting, a total of 363 HS aliquots, all derived from the same, single, homogenized human stool specimens, but prepared in four different batches, were each extracted in a different DNA extraction run. Each aliquot was processed independently on a separate DNA extraction and PCR amplification run, using the same standard operating procedure executed by a rotating group of different operators. The relative abundances of the clinical genera in each of these 363 HS aliquots were compared with each other. Overall, microbiome profiles of the HS samples were very similar to each other (Figure 4A). In contrast, a set of 400 stool samples from a subset of 897 healthy subjects described previously (Almonacid *et al.* 2017) showed very different profiles, with every subject displaying an unique pattern (Figure 4A). Beta diversity analysis showed that all HS samples clustered tightly together, irrespective of the operator, extraction robot, or sequencer (Figure 4B). However, small differences were observed between different batches of HS standards. The 363 HS replicates had an average Lin’s correlation of 0.93 (SD=0.087), which was significantly higher (KS test; p-value <0.0001) than that found among the set of 400 other stool samples, which had an average Lin’s correlation of 0.40 (SD=0.33). Lin’s correlation analysis confirmed that the effect of control batch preparation was the largest, with very limited, albeit statistically significant, effects of operator, extraction robot, or sequencer (not shown).

**Figure 4.**
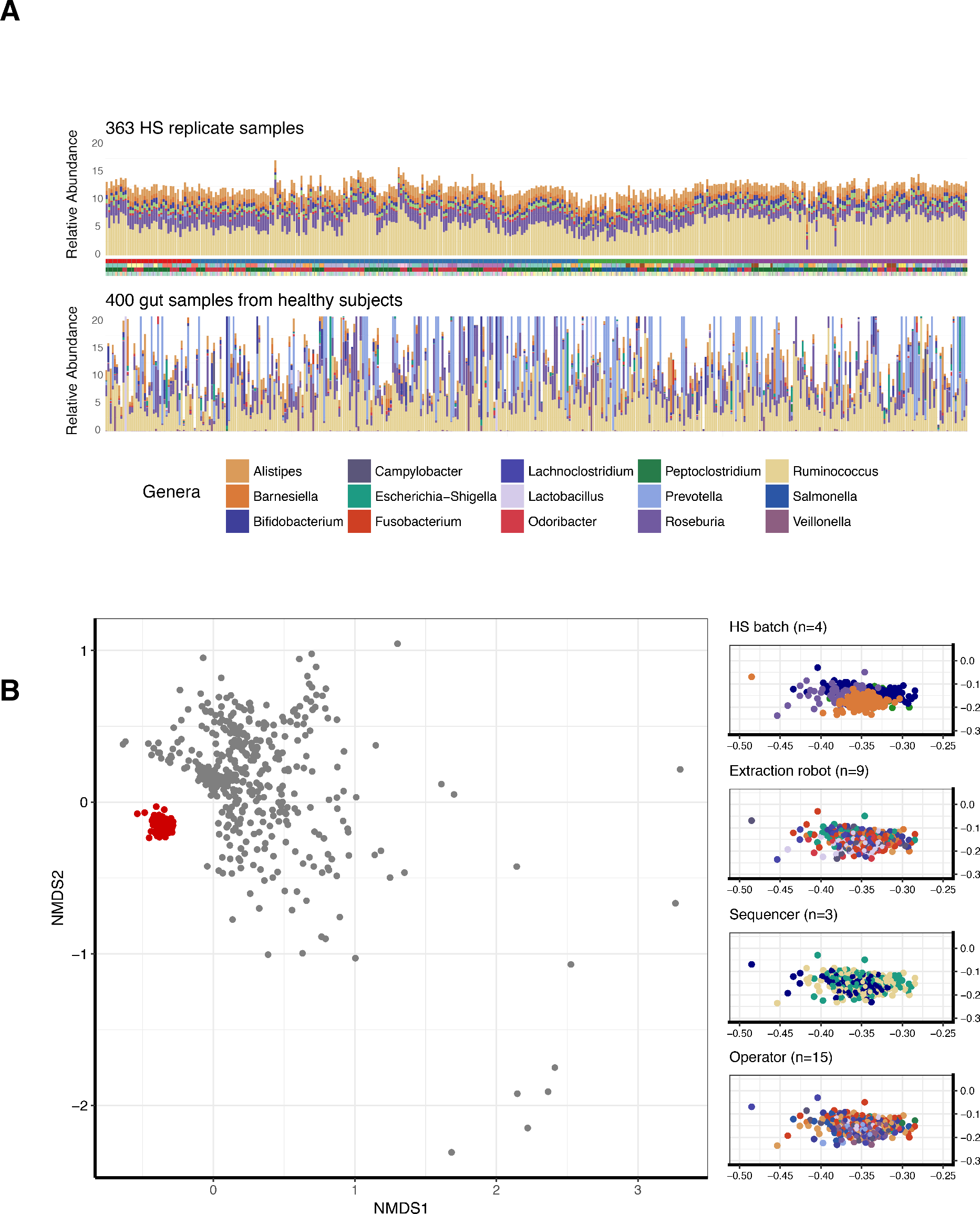
Reproducibility of microbial analysis of 363 replicate extractions of the same homogenized stool (HS) specimen. **A.** Relative abundance of clinical taxa for each of 363 HS replicates derived from subject B (top). Each HS replicate was processed in a different DNA extraction and amplification run, using different extraction robots, sequencers, and operators. Replicates are plotted in chronological order. In the bottom plot, the relative abundance of the same clinical taxa obtained from a set of 400 stool samples, each from a different subject, and none of them from subject B, is shown as a comparison. These samples were randomly selected from a group of 897 healthy subjects described in Almonacid et al. (Almonacid *et al.* 2017). **B.** Beta diversity of 363 HS replicates (in red) and the 400 other stool samples (in grey) was calculated based on the genus-level clinical taxa using Bray-Curtis dissimilarity and ordinated using NMDS. The 4 insets on the right show the ordination of the HS samples only. Each of these 4 plots shows the same data, colored according to preparation batch, sequencer machine, extraction robot, and operator, respectively.

## Discussion

Because every single step in microbial community analysis can potentially introduce biases (Pollock *et al.* 2018), a thorough understanding of the technical factors that determine variation in the outcome of microbiome analysis is essential. Choice of DNA extraction and amplification methods can have a significant impact on microbiome community patterns and their contributions are well understood (Brooks *et al.* 2015; la Cuesta-Zuluaga and Escobar 2016; Costea *et al.* 2017). However, there are several other, less understood subject and laboratory-dependent sources of variation that might cause differences in obtained microbial community profiles. Even in the setting of a high-throughput laboratory with a standardized workflow, these might contribute to unexplained variation. In this study, we investigated the effect of several of these variables.

### Homogeneity of stool

Using a set of samples taken from the same subject, we first addressed the question whether 2 pieces of stool taken from the same piece of toilet paper would be very different from each other. There is limited knowledge on how two different regions of a stool specimen have different or similar microbial compositions, but it has been argued that stool samples might not be homogeneous. Helminth egg distribution in human fecal samples, a potential measure of homogeneity, has been a topic of several studies, but these have yielded conflicting results (Krauth *et al.* 2012). Using fluorescence *in situ* hybridization and microscopy, Swidsinski and coworkers observed a differential stratification of both healthy as well as diarrheic stools (Swidsinski *et al.* 2010). Two other studies found small differences between the microbiome composition of the inner and outer areas from a fecal specimen, suggesting that subsampling of stool can result into variability of gut microbiota composition (Santiago *et al.* 2014; Gorzelak *et al.* 2015). To account for this heterogeneity and to better recover all microbes present in a fecal sample, it has been suggested that stool samples should be homogenized before DNA extraction (Gorzelak *et al.* 2015). However, homogenizing stool might not be practical for a high-throughput laboratory. Fecal material on toilet paper might actually represent a collection of different stool parts during elimination, and, in the absence of homogenisation of stool, might represent a good and practical alternative to represent the entire stool community. Although our results do not answer the question whether the microbiome of an intact stool specimen is homogenous, our study shows that taking 2 different samples from the same piece of toilet paper yield very similar microbial communities (average Lin’s correlation: 0.95).

### Reproducibility of sampling over multiple days

Another potential source of variation in microbiome composition that has not been studied extensively is how much the day-to-day variation in the diet of most humans contributes to daily changes in the gut microbiome. The same set of samples used to determine the effect of duplicate sampling from toilet paper was also used to investigate the similarity of 11 stool samples taken from the same individual within a 3- week period. Within this short time-frame, the gut microbiome of this person was relatively stable (average Lin’s correlation between all samples: 0.68), albeit significantly less similar than the paired samples obtained from the same piece of toilet paper. These results are similar to those described by others, who found there is little change in a person’s microbiome in the absence of health issues, dietary changes, or travel over a time period of days (Su *et al.* 2018) or even over longer periods (David, Materna, *et al.* 2014; Mehta *et al.* 2018). However, other studies showed that large dietary changes such as switching to a drastically different daily fiber or protein intake can change microbial composition as quickly as 2 days after starting the intervention (David, Maurice *et al.* 2014; Salonen *et al.* 2014). Together, these results suggest that microbiome profiles of healthy persons who keep a regular diet are relatively stable, while they can rapidly change as a response to large changes in diet or health status.

### Temperature stability in LS buffer

Another potential source of variation that lies outside of the internal quality control measures of a microbiome analysis laboratory is how samples are stored in the time period between obtaining the sample and their arrival in the laboratory. We wanted to test the performance of our sampling tubes, which contain a transport buffer, to preserve a stool sample for a week at 4 different temperatures, compared to a directly extracted replicate sample. There was no apparent effect of the week-long storage, even at 40°C (104°F), compared to directly extracted material (average Lin’s correlation over 8 different sample sets = 0.92). These results suggest that the proprietary buffer used in our kit is effective in preserving stool samples for a week at high temperatures. Similar results have been found with stool samples stored in ethanol or stabilization buffers such as RNAlater (Anderson *et al.* 2016; Hill *et al.* 2016; Song *et al.* 2016; Vogtman, Chen, Kibriya, *et al.* 2017; Vogtmann, Chen, Amir, *et al.* 2017; Vandeputte *et al.* 2017).

### Reproducibility of sample extraction and 16S rRNA gene microbiome analysis

Reproducibility is a critical factor in microbiome analysis. Obtaining the same results in independent rounds of DNA isolation and amplification is instrumental for the correct interpretation of results. The comparison of microbiome results obtained from replicates derived from the same sample can be used as a measure for the quality of the workflow within a laboratory. Reproducibility is important since it can be useful to evaluate quality of metagenomic or deep amplicon sequencing analyses, as seen, for example, in a metavirome study showing that replication helped to evaluate the consistency of different methods for ocean virus concentration and purification (Hurwitz, *et al.* 2013). In that study, triplicates for 3 different methods were used, being capable to observe abnormally diverse samples. Moreover, reproducibility analyses are critical for the evaluation of intra-center and inter-center performance from a standard sample (Hiergeist *et al.* 2016). In a recent paper, Raju et al. tested different sequencing protocols using 3 to 5 stool or saliva replicates and found high reproducibility in alpha diversity measures and phylum-level profiles (Raju *et al.* 2018). However, no beta diversity comparisons were provided.

Here, we report two different experiments to test the reproducibility of different parts of our laboratory workflow. In the first sample set, we tested the degree of variation introduced by three independent replicate amplification and sequencing runs performed on the same set of 44 extracted DNAs. The taxonomic composition of the three replicate analyses were very reproducible with a Lin’s correlation of 0.94. In a second experiment, we tested the reproducibility of 363 independent rounds of DNA extraction plus amplification and sequencing on the same homogenized stool sample. Even though here we added the additional variable of DNA extraction, Lin’s correlation was still excellent at 0.93. Together, these results demonstrate excellent reproducibility of different steps in our laboratory workflow, with low run to run variation. The biggest contributor in replicate sample microbiome variation was found to be the preparation of the HS control sample, which is a process that is not used for customer samples.

In the three datasets described here that include samples from different individuals, replicate samples always showed significantly higher reproducibility (Lin’s correlation over 0.9) compared to the inter-individual variation (Lin’s correlation between 0. 28 and 0.40). These results suggests that the variations introduced by replicate extractions are much smaller than the variation that is found between samples, in particular samples from different individuals.

### Standardization and quality control

Multiple calls have been made to standardize DNA extraction for microbiome marker gene or metagenomics analysis. Costea and coworkers compared 21 different DNA extraction protocols, showing big differences in DNA quality and content; they also compared different combinations of DNA isolation, library preparation and sample storage, showing the contribution of the change those methods in the variability of results; they finalize suggesting the need of a standardization for appropriate inter-lab transferability (Costea *et al.* 2017). Brooks and coworkers observed the need for 3 kinds of quality control in microbiome experiments, in order to 1) monitor the batch effects inside different sample processing runs, 2) capture variation produced by the choice of sample processing protocols and, 3) understand the difference between observed and actual community compositions for particular choices of protocols for a lab (Brooks *et al.* 2015) . In this latter work, the authors proposed the use of mixture experiments involving small “mock” communities, artificial microbial communities created by mixing known quantities of bacteria/DNA. Other studies recommended the use of careful positive (such the aforementioned mock communities or those created in chemostats) and negative controls in microbiome/metagenomic studies, in order to avoid potential pitfalls (Sinha *et al.* 2015; Kim *et al.* 2017; Pollock *et al.* 2018). Several studies also focused on the need of standardization for both feasibility and accuracy of comparative metagenomics, proposing a set of recommendations and precautions about the different strategies, including experimental and bioinformatic aspects (Knight *et al.* 2012; Nayfach and Pollard 2016).

### Final conclusions

Our results shown here show that a homogenized complete stool specimen could serve a standardized control sample in a high-throughput laboratory. The inclusion of such a sample in every extraction and amplification run, and parameters monitoring deviations from an observed microbiome profile mean, can be used to ensure the quality of the workflow. A typical complete homogenized stool specimen is enough to be aliquoted into thousands of replicates, and thus can be used to monitor thousands of microbiome analysis runs.

A limitation of this study is that the reproducible microbiome profiles shown here were obtained and analyzed using a relatively small set of microbial taxa, i.e. a small set of genus-level taxa previously reported to be associated with gastrointestinal conditions and included in our clinical assay (Almonacid *et al.* 2017). However, data analysis of subsets of the samples described here on a much larger set of genera and/or including species level taxa showed very similar reproducibility (not shown). Even with this limited set of taxa, large differences between individual stool samples could be observed.

In conclusion, we have shown that the effect of variables such as duplicate toilet paper sampling, storage temperatures, or replicate amplifications performed by different operators on different machinery is very small (Lin’s correlation over 0.9). The effect of sampling on different days was found to be somewhat larger (Lin’s correlation 0.68), but still smaller than that caused by inter-individual differences (Lin’s correlation 0.4 or lower). Including a homogenized sample into every run is an excellent way to ensure the high quality needed in a high-throughput microbiome analysis laboratory.

## Conflict of interest statement

All authors are current employees of uBiome and have received stock options and other compensation. Parts of the work described in this paper are protected by issued and provisional patents.

## Acknowledgements

Rodrigo Ortiz, Sara Bird, and Audrey Goddard are thanked for their valuable discussions and expertise. We thank all uBiome laboratory staff for their dedication, accuracy, and reproducible work.

## References

1. Al, K.F., Bisanz, J.E., Gloor, G.B., Reid, G. and Burton, J.P. 2018. Evaluation of sampling and storage procedures on preserving the community structure of stool microbiota: A simple at-home toilet-paper collection method. Journal of Microbiological Methods 144, pp. 117–121.

2. Allali, I., Arnold, J.W., Roach, J., Cadenas, M.B., Butz, N., Hassan, H.M., Koci, M., Ballou, A., Mendoza, M., Ali, R. and Azcarate-Peril, M.A. 2017. A comparison of sequencing platforms and bioinformatics pipelines for compositional analysis of the gut microbiome. BMC Microbiology 17(1), p. 194.

3. Almonacid, D.E., Kraal, L., Ossandon, F.J., Budovskaya, Y.V., Cardenas, J.P., Bik, E.M., Goddard, A.D., Richman, J. and Apte, Z.S. 2017. 16S rRNA gene sequencing and healthy reference ranges for 28 clinically relevant microbial taxa from the human gut microbiome. Plos One 12(5), p. e0176555.

4. Amir, A., McDonald, D., Navas-Molina, J.A., Debelius, J., Morton, J.T., Hyde, E., Robbins-Pianka, A. and Knight, R. 2017. Correcting for Microbial Blooms in Fecal Samples during Room-Temperature Shipping. mSystems 2(2).

5. Anderson, E.L., Li, W., Klitgord, N., Highlander, S.K., Dayrit, M., Seguritan, V., Yooseph, S., Biggs, W., Venter, J.C., Nelson, K.E. and Jones, M.B. 2016. A robust ambient temperature collection and stabilization strategy: Enabling worldwide functional studies of the human microbiome. Scientific reports 6, p. 31731.

6. Apte, Z. and Richman, J. 2015. Method and system for microbiome analysis. US Patent 9,663,831.

7. Apte, Z., Richman, J., Behbahani, S.R. and Almonacid, D. 2017. Method and system for microbiome-derived diagnostics and therapeutics. US Patent 9,703,929.

8. Bag, S., Saha, B., Mehta, O., Anbumani, D., Kumar, N., Dayal, M., Pant, A., Kumar, P., Saxena, S., Allin, K.H., Hansen, T., Arumugam, M., Vestergaard, H., Pedersen, O., Pereira, V., Abraham, P., Tripathi, R., Wadhwa, N., Bhatnagar, S., Prakash, V.G., Radha, V., Anjana, R.M., Mohan, V., Takeda, K., Kurakawa, T., Nair, G.B. and Das, B. 2016. An Improved Method for High Quality Metagenomics DNA Extraction from Human and Environmental Samples. Scientific reports 6, p. 26775.

9. Bassis, C.M., Moore, N.M., Lolans, K., Seekatz, A.M., Weinstein, R.A., Young, V.B., Hayden, M.K. and CDC Prevention Epicenters Program 2017. Comparison of stool versus rectal swab samples and storage conditions on bacterial community profiles. BMC Microbiology 17(1), p. 78.

10. Bik, E.M. 2016. The hoops, hopes, and hypes of human microbiome research. The Yale journal of biology and medicine 89(3), pp. 363–373.

11. Brooks, J.P., Edwards, D.J., Harwich, M.D., Rivera, M.C., Fettweis, J.M., Serrano, M.G., Reris, R.A., Sheth, N.U., Huang, B., Girerd, P., Vaginal Microbiome Consortium, Strauss, J.F., Jefferson, K.K. and Buck, G.A. 2015. The truth about metagenomics: quantifying and counteracting bias in 16S rRNA studies. BMC Microbiology 15, p. 66.

12. Carroll, I.M., Ringel-Kulka, T., Siddle, J.P., Klaenhammer, T.R. and Ringel, Y. 2012. Characterization of the fecal microbiota using high-throughput sequencing reveals a stable microbial community during storage. Plos One 7(10), p. e46953.

13. Choo, J.M., Leong, L.E.X. and Rogers, G.B. 2015. Sample storage conditions significantly influence faecal microbiome profiles. Scientific reports 5, p. 16350.

14. Costea, P.I., Zeller, G., Sunagawa, S., Pelletier, E., Bork, P., et al. 2017. Towards standards for human fecal sample processing in metagenomic studies. Nature Biotechnology 35(11), pp. 1069–1076.

15. de la Cuesta-Zuluaga J., and Escobar, J.S. 2016. Considerations for optimizing microbiome analysis using a marker gene. Frontiers in nutrition 3, p. 26.

16. David, L.A., Materna, A.C., Friedman, J., Campos-Baptista, M.I., Blackburn, M.C., Perrotta, A., Erdman, S.E. and Alm, E.J. 2014. Host lifestyle affects human microbiota on daily timescales. Genome Biology 15(7), p. R89.

17. David, L.A., Maurice, C.F., Carmody, R.N., Gootenberg, D.B., Button, J.E., Wolfe, B.E., Ling, A.V., Devlin, A.S., Varma, Y., Fischbach, M.A., Biddinger, S.B., Dutton, R.J. and Turnbaugh, P.J. 2014. Diet rapidly and reproducibly alters the human gut microbiome. Nature 505(7484), pp. 559–563.

18. Fischer, M.A., Güllert, S., Neulinger, S.C., Streit, W.R. and Schmitz, R.A. 2016. Evaluation of 16S rRNA Gene Primer Pairs for Monitoring Microbial Community Structures Showed High Reproducibility within and Low Comparability between Datasets Generated with Multiple Archaeal and Bacterial Primer Pairs. Frontiers in microbiology 7, p. 1297.

19. Fu, B.C., Randolph, T.W., Lim, U., Monroe, K.R., Cheng, I., Wilkens, L.R., Le Marchand, L., Hullar, M.A.J. and Lampe, J.W. 2016. Characterization of the gut microbiome in epidemiologic studies: the multiethnic cohort experience. Annals of Epidemiology 26(5), pp. 373–379.

20. Gerasimidis, K., Bertz, M., Quince, C., Brunner, K., Bruce, A., Combet, E., Calus, S., Loman, N. and Ijaz, U.Z. 2016. The effect of DNA extraction methodology on gut microbiota research applications. BMC Research Notes 9(1), p. 365.

21. Gohl, D.M., Vangay, P., Garbe, J., MacLean, A., Hauge, A., Becker, A., Gould, T.J., Clayton, J.B., Johnson, T.J., Hunter, R., Knights, D. and Beckman, K.B. 2016. Systematic improvement of amplicon marker gene methods for increased accuracy in microbiome studies. Nature Biotechnology 34(9), pp. 942–949.

22. Gorzelak, M.A., Gill, S.K., Tasnim, N., Ahmadi-Vand, Z., Jay, M. and Gibson, D.L. 2015. Methods for Improving Human Gut Microbiome Data by Reducing Variability through Sample Processing and Storage of Stool. Plos One 10(8), p. e0134802.

23. Guo, Y., Li, S.-H., Kuang, Y.-S., He, J.-R., Lu, J.-H., Luo, B.-J., Jiang, F.-J., Liu, Y.-Z., Papasian, C.J., Xia, H.-M., Deng, H.-W. and Qiu, X. 2016. Effect of shortterm room temperature storage on the microbial community in infant fecal samples. Scientific reports 6(1), p. 26648.

24. Hale, V.L., Tan, C.L., Knight, R. and Amato, K.R. 2015. Effect of preservation method on spider monkey (Ateles geoffroyi) fecal microbiota over 8 weeks. Journal of Microbiological Methods 113, pp. 16–26.

25. Hiergeist, A., Reischl, U., Priority Program 1656 Intestinal Microbiota Consortium/ quality assessment participants and Gessner, A. 2016. Multicenter quality assessment of 16S ribosomal DNA-sequencing for microbiome analyses reveals high inter-center variability. International Journal of Medical Microbiology 306(5), pp. 334–342.

26. Hill, C.J., Brown, J.R.M., Lynch, D.B., Jeffery, I.B., Ryan, C.A., Ross, R.P., Stanton, C. and O’Toole, P.W. 2016. Effect of room temperature transport vials on DNA quality and phylogenetic composition of faecal microbiota of elderly adults and infants. Microbiome 4(1), p. 19.

27. Hurwitz, B.L., Deng, L., Poulos, B.T. and Sullivan, M.B. 2013. Evaluation of methods to concentrate and purify ocean virus communities through comparative, replicated metagenomics. Environmental Microbiology 15(5), pp. 1428–1440.

28. Jones, M.B., Highlander, S.K., Anderson, E.L., Li, W., Dayrit, M., Klitgord, N., Fabani, M.M., Seguritan, V., Green, J., Pride, D.T., Yooseph, S., Biggs, W., Nelson, K.E. and Venter, J.C. 2015. Library preparation methodology can influence genomic and functional predictions in human microbiome research. Proceedings of the National Academy of Sciences of the United States of America 112(45), pp. 14024–14029.

29. Jovel, J., Patterson, J., Wang, W., Hotte, N., O’Keefe, S., Mitchel, T., Perry, T., Kao, D., Mason, A.L., Madsen, K.L. and Wong, G.K.-S. 2016. Characterization of the gut microbiome using 16S or shotgun metagenomics. Frontiers in microbiology 7, p. 459.

30. Kennedy, N.A., Walker, A.W., Berry, S.H., Duncan, S.H., Farquarson, F.M., Louis, P., Thomson, J.M., UK IBD Genetics Consortium, Satsangi, J., Flint, H.J., Parkhill, J., Lees, C.W. and Hold, G.L. 2014. The impact of different DNA extraction kits and laboratories upon the assessment of human gut microbiota composition by 16S rRNA gene sequencing. Plos One 9(2), p. e88982.

31. Kim, D., Hofstaedter, C.E., Zhao, C., Mattei, L., Tanes, C., Clarke, E., Lauder, A., Sherrill-Mix, S., Chehoud, C., Kelsen, J., Conrad, M., Collman, R.G., Baldassano, R., Bushman, F.D. and Bittinger, K. 2017. Optimizing methods and dodging pitfalls in microbiome research. Microbiome 5(1), p. 52.

32. Klindworth, A., Pruesse, E., Schweer, T., Peplies, J., Quast, C., Horn, M. and Glöckner, F.O. 2013. Evaluation of general 16S ribosomal RNA gene PCR primers for classical and next-generation sequencing-based diversity studies. Nucleic Acids Research 41(1), p. e1.

33. Knight, R., Callewaert, C., Marotz, C., Hyde, E.R., Debelius, J.W., McDonald, D. and Sogin, M.L. 2017. The microbiome and human biology. Annual Review of Genomics and Human Genetics 18, pp. 65–86.

34. Knight, R., Jansson, J., Field, D., Fierer, N., Desai, N., Fuhrman, J.A., Hugenholtz, P., van der Lelie, D., Meyer, F., Stevens, R., Bailey, M.J., Gordon, J.I., Kowalchuk, G.A. and Gilbert, J.A. 2012. Unlocking the potential of metagenomics through replicated experimental design. Nature Biotechnology 30(6), pp. 513–520.

35. Knudsen, B.E., Bergmark, L., Munk, P., Lukjancenko, O., Priemé, A., Aarestrup, F.M. and Pamp, S.J. 2016. Impact of sample type and DNA isolation procedure on genomic inference of microbiome composition. mSystems 1(5).

36. Krauth, S.J., Coulibaly, J.T., Knopp, S., Traoré, M., N’Goran, E.K. and Utzinger, J. 2012. An in-depth analysis of a piece of shit: distribution of Schistosoma mansoni and hookworm eggs in human stool. PLoS Neglected Tropical Diseases 6(12), p. e1969.

37. Letunic, I. and Bork, P. 2016. Interactive tree of life (iTOL) v3: an online tool for the display and annotation of phylogenetic and other trees. Nucleic Acids Research 44(W1), pp. W242–5.

38. Lin, L.I. 1989. A concordance correlation coefficient to evaluate reproducibility. Biometrics 45(1), pp. 255–268.

39. Mahé, F., Rognes, T., Quince, C., de Vargas, C. and Dunthorn, M. 2014. Swarm: robust and fast clustering method for amplicon-based studies. PeerJ 2, p. e593.

40. Massey, F.J. 1951. The Kolmogorov-Smirnov Test for Goodness of Fit. Journal of the American Statistical Association 46(253), pp. 68–78.

41. McMurdie, P.J. and Holmes, S. 2013. phyloseq: an R package for reproducible interactive analysis and graphics of microbiome census data. Plos One 8(4), p. e61217.

42. Mehta, R.S., Abu-Ali, G.S., Drew, D.A., Lloyd-Price, J., Subramanian, A., Lochhead, P., Joshi, A.D., Ivey, K.L., Khalili, H., Brown, G.T., DuLong, C., Song, M., Nguyen, L.H., Mallick, H., Rimm, E.B., Izard, J., Huttenhower, C. and Chan, A.T. 2018. Stability of the human faecal microbiome in a cohort of adult men. Nature microbiology 3(3), pp. 347–355.

43. Nature Microbiology Editorial 2016. Raising standards in microbiome research. Nature microbiology 1(7), p. 16112.

44. Nayfach, S. and Pollard, K.S. 2016. Toward accurate and quantitative comparative metagenomics. Cell 166(5), pp. 1103–1116.

45. Oksanen, J., Blanchet, F.G., Friendly, M., Kindt, R., Legendre, P., McGlinn, D., Minchin, P.R., O’Hara, R.B., Simpson, G.L., Solymos, P., Stevens, M.H.H., Szoecs, E. and Wagner, H. 2016. vegan: Community Ecology Package. R package version 2.4-1. [Online]. Available at: https://CRAN.R-project.org/package=vegan.

46. Pollock, J., Glendinning, L., Wisedchanwet, T. and Watson, M. 2018. The madness of microbiome: Attempting to find consensus “best practice” for 16S microbiome studies. Applied and Environmental Microbiology, p. AEM.02627–17.

47. Quast, C., Pruesse, E., Yilmaz, P., Gerken, J., Schweer, T., Yarza, P., Peplies, J. and Glöckner, F.O. 2013. The SILVA ribosomal RNA gene database project: improved data processing and web-based tools. Nucleic Acids Research 41(Database issue), pp. D590–6.

48. Raju, S.C., Lagström, S., Ellonen, P., de Vos, W.M., Eriksson, J.G., Weiderpass, E. and Rounge, T.B. 2018. Reproducibility and repeatability of six high throughput 16S rDNA sequencing protocols for microbiota profiling. Journal of Microbiological Methods 147, pp. 76–86.

49. Ranjan, R., Rani, A., Metwally, A., McGee, H.S. and Perkins, D.L. 2016. Analysis of the microbiome: Advantages of whole genome shotgun versus 16S amplicon sequencing. Biochemical and Biophysical Research Communications 469(4), pp. 967–977.

50. Röder, B., Frühwirth, K., Vogl, C., Wagner, M. and Rossmanith, P. 2010. Impact of long-term storage on stability of standard DNA for nucleic acid-based methods. Journal of Clinical Microbiology 48(11), pp. 4260–4262.

51. Rognes, T., Flouri, T., Nichols, B., Quince, C. and Mahé, F. 2016. VSEARCH: a versatile open source tool for metagenomics. PeerJ 4, p. e2584.

52. Romanazzi, V., Traversi, D., Lorenzi, E. and Gilli, G. 2015. Effects of freezing storage on the DNA extraction and microbial evaluation from anaerobic digested sludges. BMC Research Notes 8, p. 420.

53. Salonen, A., Lahti, L., Salojärvi, J., Holtrop, G., Korpela, K., Duncan, S.H., Date, P., Farquharson, F., Johnstone, A.M., Lobley, G.E., Louis, P., Flint, H.J. and de Vos, W.M. 2014. Impact of diet and individual variation on intestinal microbiota composition and fermentation products in obese men. The ISME Journal 8(11), pp. 2218–2230.

54. Santiago, A., Panda, S., Mengels, G., Martinez, X., Azpiroz, F., Dore, J., Guarner, F. and Manichanh, C. 2014. Processing faecal samples: a step forward for standards in microbial community analysis. BMC Microbiology 14, p. 112.

55. Schnorr, S.L., Candela, M., Rampelli, S., Centanni, M., Consolandi, C., Basaglia, G., Turroni, S., Biagi, E., Peano, C., Severgnini, M., Fiori, J., Gotti, R., De Bellis, G., Luiselli, D., Brigidi, P., Mabulla, A., Marlowe, F., Henry, A.G. and Crittenden, A.N. 2014. Gut microbiome of the Hadza hunter-gatherers. Nature Communications 5(5), p. 3654.

56. Schultze, A., Akmatov, M.K., Andrzejak, M., Karras, N., Kemmling, Y., Maulhardt, A., Wieghold, S., Ahrens, W., Günther, K., Schlenz, H., Krause, G. and Pessler, F. 2014. Comparison of stool collection on site versus at home in a population-based study: feasibility and participants’ preference in Pretest 2 of the German National Cohort. Bundesgesundheitsblatt, Gesundheitsforschung, Gesundheitsschutz 57(11), pp. 1264–1269.

57. Sinha, R., Abnet, C.C., White, O., Knight, R. and Huttenhower, C. 2015. The microbiome quality control project: baseline study design and future directions. Genome Biology 16, p. 276.

58. Sinha, R., Abu-Ali, G., Vogtmann, E., Fodor, A.A., Ren, B., Amir, A., Schwager, E., Crabtree, J., Ma, S., Microbiome Quality Control Project Consortium, Abnet, C.C., Knight, R., White, O. and Huttenhower, C. 2017. Assessment of variation in microbial community amplicon sequencing by the Microbiome Quality Control (MBQC) project consortium. Nature Biotechnology 35(11), pp. 1077–1086.

59. Sinha, R., Chen, J., Amir, A., Vogtmann, E., Shi, J., Inman, K.S., Flores, R., Sampson, J., Knight, R. and Chia, N. 2016. Collecting fecal samples for microbiome analyses in epidemiology studies. Cancer Epidemiology, Biomarkers & Prevention 25(2), pp. 407–416.

60. Song, S.J., Amir, A., Metcalf, J.L., Amato, K.R., Xu, Z.Z., Humphrey, G. and Knight, R. 2016. Preservation methods differ in fecal microbiome stability, affecting suitability for field studies. mSystems 1(3).

61. Su, T., Liu, R., Long, Y., Quan, S., Lai, S., Wang, L., Si, J. and Chen, S. 2018. 1-Day or 5-Day Fecal Samples, Which One is More Beneficial to be Used for DNA Based Gut Microbiota Study. Current Microbiology 75(3), pp. 288–295.

62. Swidsinski, A., Loening-Baucke, V., Kirsch, S. and Doerffel, Y. 2010. Functional biostructure of colonic microbiota (central fermenting area, germinal stock area and separating mucus layer) in healthy subjects and patients with diarrhea treated with Saccharomyces boulardii. Gastroentérologie Clinique et Biologique 34, pp. S79–S92.

63. Tedjo, D.I., Jonkers, D.M.A.E., Savelkoul, P.H., Masclee, A.A., van Best, N., Pierik, M.J. and Penders, J. 2015. The effect of sampling and storage on the fecal microbiota composition in healthy and diseased subjects. Plos One 10(5), p. e0126685.

64. Tessler, M., Neumann, J.S., Afshinnekoo, E., Pineda, M., Hersch, R., Velho, L.F.M., Segovia, B.T., Lansac-Toha, F.A., Lemke, M., DeSalle, R., Mason, C.E. and Brugler, M.R. 2017. Large-scale differences in microbial biodiversity discovery between 16S amplicon and shotgun sequencing. Scientific reports 7(1), p. 6589.

65. Thatcher, S.A. 2015. DNA/RNA preparation for molecular detection. Clinical Chemistry 61(1), pp. 89–99.

66. Thijs, S., Op De Beeck, M., Beckers, B., Truyens, S., Stevens, V., Van Hamme, J.D., Weyens, N. and Vangronsveld, J. 2017. Comparative Evaluation of Four Bacteria-Specific Primer Pairs for 16S rRNA Gene Surveys. Frontiers in microbiology 8, p. 494.

67. Thomas, V., Clark, J. and Doré, J. 2015. Fecal microbiota analysis: an overview of sample collection methods and sequencing strategies. Future Microbiology 10(9), pp. 1485–1504.

68. Tyler, A.D., Smith, M.I. and Silverberg, M.S. 2014. Analyzing the human microbiome: a “how to” guide for physicians. The American Journal of Gastroenterology 109(7), pp. 983–993.

69. Vandeputte, D., Tito, R.Y., Vanleeuwen, R., Falony, G. and Raes, J. 2017. Practical considerations for large-scale gut microbiome studies. FEMS Microbiology Reviews 41(Supp_1), pp. S154–S167.

70. Vogtmann, E., Chen, J., Amir, A., Shi, J., Abnet, C.C., Nelson, H., Knight, R., Chia, N. and Sinha, R. 2017. Comparison of collection methods for fecal samples in microbiome studies. American Journal of Epidemiology 185(2), pp. 115–123.

71. Vogtmann, E., Chen, J., Kibriya, M.G., Chen, Y., Islam, T., Eunes, M., Ahmed, A., Naher, J., Rahman, A., Amir, A., Shi, J., Abnet, C.C., Nelson, H., Knight, R., Chia, N., Ahsan, H. and Sinha, R. 2017. Comparison of fecal collection methods for microbiota studies in Bangladesh. Applied and Environmental Microbiology 83(10).

72. Walker, A.W., Martin, J.C., Scott, P., Parkhill, J., Flint, H.J. and Scott, K.P. 2015. 16S rRNA gene-based profiling of the human infant gut microbiota is strongly influenced by sample processing and PCR primer choice. Microbiome 3(1), p. 26.

73. Ward, J.H. 1963. Hierarchical Grouping to Optimize an Objective Function. Journal of the American Statistical Association 58(301), pp. 236–244.

74. Wong, W.S.W., Clemency, N., Klein, E., Provenzano, M., Iyer, R., Niederhuber, J.E. and Hourigan, S.K. 2017. Collection of non-meconium stool on fecal occult blood cards is an effective method for fecal microbiota studies in infants. Microbiome 5(1), p. 114.

75. Young, V.B. 2017. The role of the microbiome in human health and disease: an introduction for clinicians. BMJ.

76. Yuan, S., Cohen, D.B., Ravel, J., Abdo, Z. and Forney, L.J. 2012. Evaluation of methods for the extraction and purification of DNA from the human microbiome. Plos One 7(3), p. e33865.

